# Ultrastructure Expansion Microscopy (U-ExM) in *Trypanosoma cruzi*: localization of tubulin isoforms and isotypes

**DOI:** 10.1101/2022.03.06.483067

**Authors:** Victoria Lucia Alonso

## Abstract

Ultrastructure Expansion Microscopy (U-ExM) is a recently developed technique that enables the increase of the spatial resolution within a cell or a tissue for microscopic imaging by physically expanding the sample. For the first time, I report a detailed protocol validating the use of U-ExM in *Trypanosoma cruzi* and quantifying the expansion factors of different subcellular compartments. I was able to determine the localization patterns of different tubulin isoforms, such as α-tubulin and β-tubulin. Also, I immunolocalized acetylated and tyrosinated α-tubulin isotypes in epimastigotes and use mitochondrial cell-permeable dyes to identify this organelle. Finally, U-ExM was also performed in trypomastigotes and amastigotes validating this technique in all life cycle stages of *T. cruzi*.

## Introduction

*Trypanosoma cruzi*, a kinetoplastid parasite and the etiological agent of Chagas disease or American trypanosomiasis, has with a complex life cycle that alternates between a mammalian and an insect host (Triatominidae family), which is the biological vector of this disease (Lidani et al. 2019). The World Health Organization classifies Chagas disease as one of the 13 most neglected tropical diseases, constituting a very important social and economic problem, especially in Latin America.

Trypanosomatids are characterized by a precisely organized cytoskeletal constituted by stable microtubules (MT), which is simpler in comparison with other eukaryotic cells. Such MTs are present in the mitotic spindle, the basal body, the flagellar axoneme, and the subpellicular microtubules. The latter are connected to each other and to the plasma membrane forming a helical arrangement along the central axis of the parasite cell body (de Souza 2009). MTs are composed by α/β-tubulin heterodimers forming helical tubes and provide the basis for cytoskeletal architecture. They are regulated by interacting with a variety of MT-associated proteins (MAPs), by a differential expression of α- and β-tubulin genes (tubulin isotypes), and by tubulin isoforms generated by the inclusion of different post-translational modifications (PTMs) in each isotype (Gadadhar et al. 2017). In trypanosomatids, many PTMs were identified across the microtubular array. It is proposed that this could be used to control the differential recruitment of MAPs and motors during the cell cycle (Sinclair and de Graffenried 2019).

Ultrastructure Expansion microscopy (U-ExM) has been introduced recently as a super-resolution microscopy technique. Basically, biological specimens are included in swellable polymer hydrogels and then are physically expanded to resolve fine details (Chen et al. 2015). This method its very simple and accessible, its compatibility with conventional fluorescent probes, and inexpensive chemicals. It can achieve a spatial resolution of ~65 nm using conventional (confocal) microscopes, around four times better than the standard resolution of a confocal microscope (~250 nm).

Since 2015 there has been a steady growth of the publication involving this technique, especially in protozoan parasites, like *Giardia lamblia, Plasmodium falciparum*, and *Toxoplasma gondii* (Tillberg and Chen 2019; dos Santos Pacheco and Soldati-Favre 2021; Gorilak et al. 2021; Tomasina et al. 2021). This technique was first reported in the kinetoplastid *Trypanosoma brucei*, where a component of the mitochondrial genome segregation machinery was described (Amodeo et al. 2021; Kalichava and Ochsenreiter 2021), but it has not been reported in *T. cruzi*.

In this report, I validate the use of U-ExM in *Trypanosoma cruzi* and quantify the expansion factors of different subcellular structures. I describe the localization patterns of tubulin isoforms, such as α-tubulin, β-tubulin, validating this technique in all life cycle stages of *T. cruzi*. Also, I immunolocalized acetylated and tyrosinated α-tubulin isotypes in epimastigotes to visualize subpellicular microtubules, the flagellar axoneme, the basal body, and the flagellar pocket. Furthermore, I validated the use of the cell-permeable dyes, such as Mitotracker, for U-ExM in *T. cruzi*.

## Methods

### Parasite culture and infections

*T. cruzi* Dm28*c* epimastigotes and Vero cells (ATCC CCL-81) were cultured and infected as previously described (Alonso et al. 2016).

### U-ExM protocol

I started with 5 million epimastigotes or trypomastigotes, washed the pellets twice with 500 μl PBS and centrifuged for 5 min at 2000 g, then resuspended in 200 μl of PBS and settled for 20 min at room temperature on poly-D-lysine coated coverslips (12 mm, Marlenfeld GmbH). For mitochondrial staining, epimastigotes were resuspended in PBS and incubated with 1mM Mitotracker Orange CMTMRos (Invitrogen) for 30 min at 28°C, washed twice in PBS and fixed with 3% paraformaldehyde + 0.1% Glutaraldehyde (Sigma, 70% solution) in PBS for 15 min at RT and then settled on coverslips as described before. Cells infected with amastigotes in 12 mm coverslips were washed twice with PBS to eliminate culturing media. For all the conditions stated above, coverslips were carefully submerged into a 24-well plate with 1 ml of 4% Formaldehyde (FA, 36.5–38%, Sigma) and 4% acrylamide (AA, 40% stock solution, Invitrogen) in PBS and incubated for four to five hours at 37°C. For gelation, coverslips were putted, with the cells facing down, into 35μl of monomer solution: 19% Sodium Acrylate (Sigma), 10% AA, 0.1% N, N’-methylenbisacrylamide (BIS, Sigma) in PBS, supplemented with 0.5% APS (Sigma) and 0.5% tetramethylethylenediamine (TEMED, Sigma). A drop of monomer solution was placed over a parafilm in a pre-cooled humid chamber for each coverslip. For gelation, samples were incubated 5 mins on ice, and then for 1 h at 37°C in the dark. Coverslips with gels were then transferred into a six-well plate filled with 2 ml of denaturation buffer (200 mM sodium dodecyl sulfate (SDS), 200 mM NaCl, and 50 mM Tris-HCl in MilliQ, pH 9) for 15 min at room temperature with gentle agitation to allow the gels to detach from the coverslips. Gels were then moved into a 1.5 ml Eppendorf centrifuge tube filled with 1 ml denaturation buffer and incubated at 95°C for 1 h and 30 min. After denaturation, gels were placed in 50 ml beakers or Petri dishes filled with MilliQ water for the first round of expansion. Water was exchanged after 30 minutes and then gels were incubated overnight in MilliQ water. The next day, water was exchanged one more time for 30 minutes and then gels were washed two times for 30 min in PBS (to allow gel shrinkage) and subsequently incubated with primary antibody (Table I), all antibodies were diluted in 2% PBS/BSA, and gels were incubated for 3 h at 37°C and then overnight at 4°C in 24 well plates with 300 μl of the diluted antibodies. Gels were then washed in PBS-T three times for 10 min while gently shaking and subsequently incubated with secondary antibodies (Table I) using the same incubation times. The day after, gels were washed in PBS-T (three times for 10 min while shaking) and placed into beakers or Petri dishes filled with MilliQ water for the second round of expansion. Water was exchanged three times every 30 min. After this step, gels were carefully measured with a caliper showing an expansion factor between 4.2x and 4.6x. The specificity of the antibodies was verified by western blot with epimastigotes’ total extracts (Figure 1a).

**Fig. 1.**
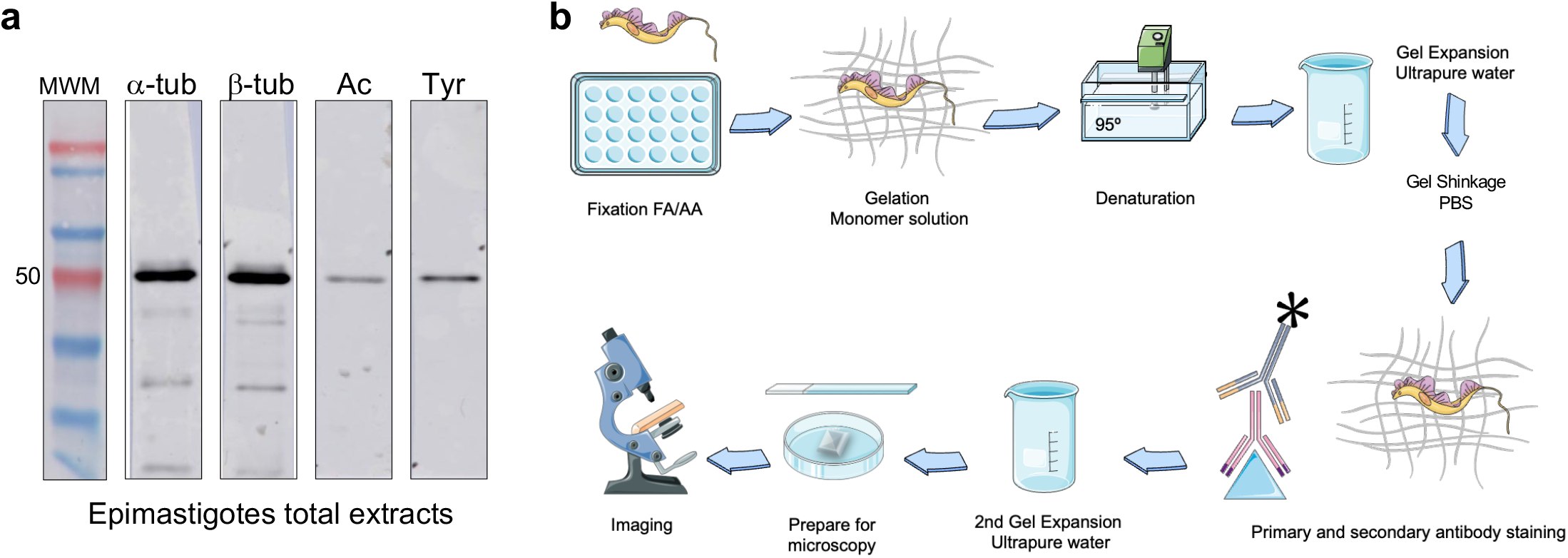
Specificity of the anti-α-tubulin (49,8 kDa), β-tubulin (49.7 kDa), acetylated (Ac) α-tubulin and tyrosinated (Tyr) α-tubulin antibodies were verified by western blot against total epimastigotes extracts. 1 million parasites were blotted per lane, goat anti-mouse IgG conjugated to HRP was used as the secondary antibody. **b)** U-ExM workflow in *T. cruzi*. Servier Medical Art was used to create this figure (smart.servier.com)

### Imaging and measurement of the expansion

The gel was then cut with a razor blade into pieces and put into a 35 mm glass-bottom dish (Matsunami) previously coated with poly-lysine for confocal imaging. Confocal microscopy was performed on a Zeiss LSM880 using a 63x oil NA 1.4 objective, with the following parameters: z step size at 0.65 μm interval with a pixel size of 0.132 nm. ZEN BLACK 2.1 SP3 software (Zeiss, RRID:SCR_018163) and Fiji (Schindelin et al. 2012) (RRID:SCR_002285) were used to analyze the images.

To determine the expansion factor, the ratio of expanded kinetoplast and nucleus to non-expanded epimastigotes was calculated (n = 20 cells) using GraphPad Prism v. 9.0.1 (RRID:SCR_002798). The length of the kinetoplast was determined by measuring its maximum length and the diameter of the nucleus by measuring the widest diameter of each epimastigote using Fiji.

## Results and Discussion

### U-ExM workflow in *T. cruzi*

Although the Ex-M protocols present in the literature follow the same steps, they have differences regarding the concentration of fixative agents, denaturing agents, and incubation times (this also depends on the nature of the protein to be immunolocalized). I tested several conditions and optimized the workflow for *T. cruzi* (Figure 1), which is described in Methods section. In this report, I used antibodies against cytoskeletal proteins (insoluble proteins) (Table I) and gave a detailed protocol to perform the U-ExM, but is important to mention that the incubation time when targeting soluble proteins or membrane proteins the incubation time and temperature during the denaturing step should be optimized.

As observed in Figure 1, in the first step, FA and AA (precursor molecules) are introduced to functionalize the cellular components, this allows the subsequent linkage of swellable polyelectrolyte gel (Monomer solution containing SA). The gel formation is accomplished using the oxidizing reagents TEMED and APS (as for standard polyacrylamide gels). Then, the parasite contents are denatured using SDS and high temperature. After this step, the first round of expansion is performed in ultrapure water followed by incubation in PBS, where the gel shrinks again. Finally, standard immunofluorescence techniques are applied followed by the second round of expansion in water before imaging in an epifluorescence or confocal microscope.

### Localization of tubulin isoforms

MTs are essential components of the eukaryotic cytoskeleton and cell division machinery. I immunolocalized α- and β-tubulina using specific monoclonal antibodies (Figures 2a and 2b). A similar localization pattern was observed for both isoforms at the subpellicular microtubules and the flagellar axoneme. The axoneme is immunostained from the base of the flagellum to its tip, in continuous attachment to the cell body as previously described (de Souza 2009)). Pro-and mature basal bodies were also clearly observed, as well as the microtubule quartet in proximity to the kinetoplast in a parasite that is starting to divide (Zoom in Figure 2b) (Elias et al. 2007).

**Fig. 2.**
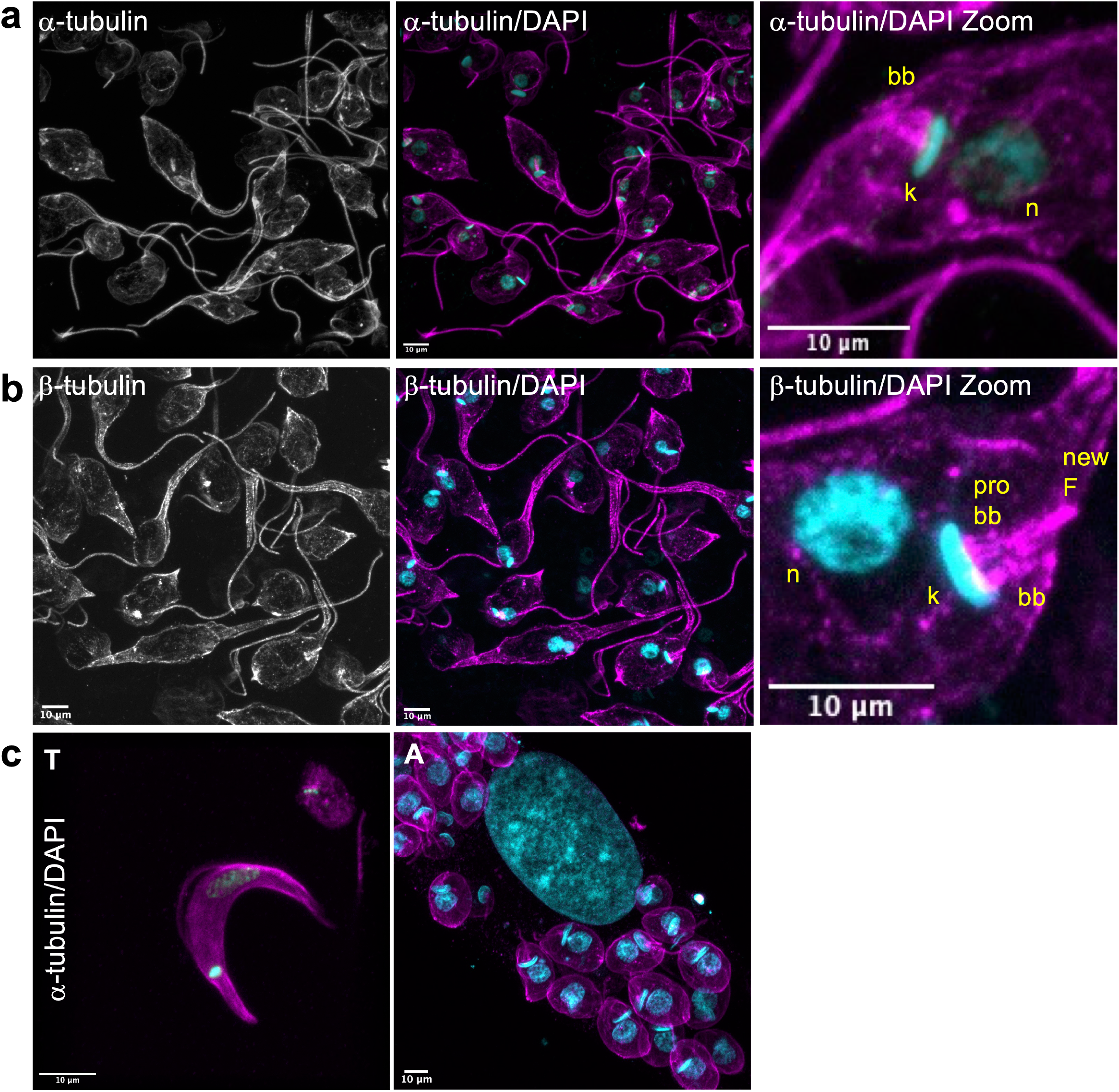
Tubulin isotypes during *T. cruzi* cell stages. **a)** Epimastigotes stained with anti α-tubulin antibodies (greyscale or magenta; Alexa 555) and DAPI (cyan; kDNA and nuclear DNA). In Zoom images: bb, basal body; k, linetoplast; n, nucleus **b)** Epimastigotes stained with anti β-tubulin antibodies (greyscale or magenta; Alexa 555) and DAPI (cyan; kDNA and nuclear DNA). In Zoom images: bb, basal body; pro-bb, pro basal body; k, kinetoplast; n, nucleus, F, flagella **c)** Trypomastogotes (T) and amastigotes (A) stained with anti α-tubulin antibodies (magenta; Alexa 555) and DAPI (cyan; kDNA and nuclear DNA).

Alpha-tubulin was also immunolocalized in trypomastigotes and amastigotes, highlighting this technique’s value in all life cycle stages of *T. cruzi*. As observed in figure 2c the subpellicular array and the flagellar axoneme are clearly immunostained in both stages. It is interesting to note that the anti α-tubulin antibody used (TAT-1), immunodetects, in our experimental conditions, exclusively amastigotes’α-tubulin and not the host cell protein, I believe this is because TAT-1 was developed by immunizing mice with extracted *T. brucei brucei* cytoskeleton although it is suitable to use in mammalian cells.

### Localization of tubulin isotypes in *T. cruzi*

There are many conserved acetylated lysines (K) reported in α- and β-tubulin, but acetylation of the α-tubulin luminal residue K40, discovered over thirty years ago, has been widely characterized. This acetylation mark is associated with stable microtubes and with the resistant microtubules to depolymerizing drugs (L’Hemault and Rosenbaum 1985). In Trypanosomatids subpellicular, mitotic and axonemal microtubules are extensively acetylated making them interesting models for studying K40 acetylation (Sasse and Gull 1988; Souto-Padron et al. 1993). As observed in Figure 3, acetylated α-tubulin is presenten in the subpellicular microtubules and flagellar axoneme. Immunodetection of the flagellar pocket (Figure 3a) and the basal body is also observed in great detail using U-ExM. Tyrosinated α-tubulin shows a different localization pattern (Figure 3b). The subpellicular MTs of the posterior end of the cell body, the nascent flagellum of dividing cells, and the basal bodies are all immunodetected, similar to the localization reported for *T. brucei* (Sherwin et al. 1987). A weak reaction is detected in the subpellicular MTs of the anterior part of the epimastigotes and the old flagellum.

**Fig. 3.**
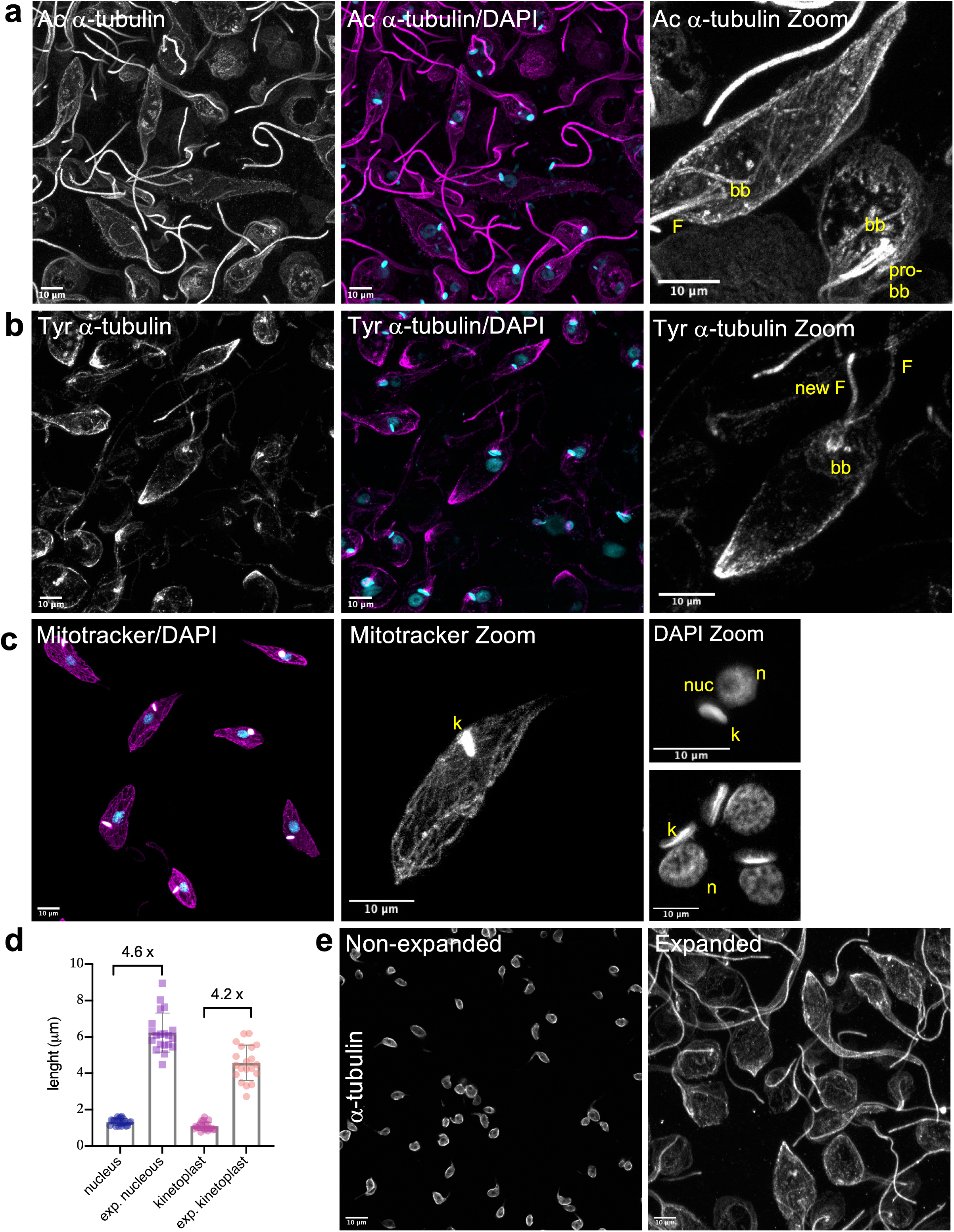
Tubulin isoforms, mitochondrial staining and expansion measurments in epimastigotes. **a)** Epimastigotes stained with anti Acetylated (Ac) α-tubulin antibodies (greyscale or magenta; Alexa 555) and DAPI (cyan; kDNA and nuclear DNA). In Zoom images: bb, basal body; F, flagella **b)** Epimastigotes stained with anti Tyrosonated (Tyr) α-tubulin antibodies (greyscale or magenta; Alexa 555) and DAPI (cyan; kDNA and nuclear DNA). In Zoom images: bb, basal body; F, flagella **c)** Epimastigotes stained with the cell-permeable dye Mitotracker and DAPI. In Zoom images: k, kinetoplast; n, nucleus; nuc, nucleolus **d)** Nucleus and kinetoplast measurements in expanded and non-expanded epimastigotes using Fiji. Expansion factor was calculated as the ratio between non-expanded and expanded measurements **e)** Non-expanded and expanded epimastigotes stained with anti α-tubulin (greyscale; Alexa 555).

### Measurement of subcellular compartments and structures in epimastigotes

Trypanosomatids are characterized by a single flagellum and a mass of mitochondrial DNA organized into a kinetoplast connected to the proximal end of the basal bodies and the flagellar pocket (Gull 1999). The kinetoplast was measured in epimastigotes stained with Mitotracker and for nuclear measurements, I stained epimastigotes’ DNA with DAPI (Figure 3c). With Mitotracker staining, the ramifications of the single mitochondria of *T. cruzi* are clearly identified and the kinetoplast is deeply marked, this also allowed us to validate the use of cell-permeable dyes for U-ExM in *T. cruzi*. Using DAPI I was able to discriminate condensed regions of the chromatin as well as the nucleolar domain (identified by the lack of DAPI staining) (Figure 3c). An expansion factor of 4.2x and 4.6x was quantified for the nucleus and the kinetoplast respectively, which signifies that expansion was homogeneous within the cells (Figure 3d).

When compared with non-expanded epimastigotes, the resolution gain is clear and allowed us to visualize the potential of this technique in *T. cruzi* (Figure 3e). In conclusion, U-ExM is a powerful approach for studying cytoskeletal structures in *T. cruzi*. It is a robust super-resolution method, but it is very simple and does not need highly specialized reagents or instrumentation. U-ExM allows high-resolution imaging, improving the axial and lateral effective resolution of the molecules to be analyzed. An expansion factor of four increases resolution from around 250 nm to less than 70 nm using regular confocal microscopy. In the future, this could be further improved by super-resolution techniques to visualize biomolecular structures that are in close proximity to each other, such as the components of the flagellar pocket and basal body, which are poorly described in *T. cruzi*.

## Statements and Declarations

### Competing interest

The authors have no relevant financial or non-financial interests to disclose.

### Funding

This work was supported by Agencia Nacional de Promoción Cientifica y Tecnológica from Argentina (PICT2017-1978 and PICT2019-0526) and the Research Council United Kingdom (MR/P027989/1).

### Authors contribution statement

VLA has designed, performed the experiments, and wrote the manuscript.

## Acknowledgments

Thanks to Dr. Esteban Serra and Dr. Luis E. Tavernelli for the critical reading of the manuscript. I am also very grateful to Rodrigo Vena for assisting in the acquisition of confocal images.

**Table I: List of antibodies used and working dilutions**

## Notes

### Competing Interest Statement

The authors have declared no competing interest.

### Summary of Updates

Manuscript was updated to clarify.

